# Identification of a binding site for bupropion in *Gloeobacter violaceus* ligand-gated ion channel

**DOI:** 10.1101/2023.10.09.561596

**Authors:** Elham Pirayesh, Hoa Quynh Do, Garren Ferreira, Akash Pandhare, Zackary Ryan Gallardo, Michaela Jansen

**Affiliations:** Medical Student Summer Research Program, School of Medicine, Texas Tech University Health Sciences Center, Lubbock, TX, 79430 USA; Cell Physiology Molecular Biophysics, School of Medicine, Texas Tech University Health Sciences Center, Lubbock, TX, 79430 USA

## Abstract

Bupropion is an atypical antidepressant and smoking cessation drug which causes adverse effects such as insomnia, irritability, and anxiety. Bupropion inhibits dopamine and norepinephrine reuptake transporters and eukaryotic cation-conducting pentameric ligand-gated ion channels (pLGICs), such as nicotinic acetylcholine (nACh) and serotonin type 3A (5-HT3A) receptors, at clinically relevant concentrations. However, the binding sites and binding mechanisms of bupropion are still elusive. To further understand the inhibition of pLGICs by bupropion, in this work, using a prokaryotic homologue of pLGICs as a model, we examined the inhibitory potency of bupropion in *Gloeobacter violaceus* ligand-gated ion channel (GLIC), a proton-gated ion channel. Bupropion inhibited proton-induced currents in GLIC with an inhibitory potency of 14.9 ± 2.0 μM, comparable to clinically attainable concentrations previously shown to also modulate eukaryotic pLGICs. Using single amino acid substitutions in GLIC and two-electrode voltage-clamp recordings, we further determined a binding site for bupropion in the lower third of the first transmembrane segment M1 at residue T214. The sidechain of M1 T214 together with additional residues of M1 and also of M3 of the adjacent subunit have previously been shown to contribute to binding of other lipophilic molecules like allopregnanolone and pregnanolone.

**SIGNIFICANCE:** GLIC, *Gloeobacter* ligand-gated ion channel, has been extensively used as a model to identify and understand binding sites and mechanisms for cholesterol, neurosteroids, and anesthetics interacting with neurotransmitter-gated ion channels, such as, GABA_A_ receptors and nicotinic acetylcholine receptors. Recently, increasing evidence has revealed that another neurotransmitter-gated ion channel, the serotonin type 3A receptor, binds to an atypical antidepressant, bupropion, at clinically relevant concentrations. Our work here proposes a binding site for bupropion in GLIC and suggests a new approach to characterize binding mechanisms of bupropion in other neurotransmitter-gated ion channels.

## INTRODUCTION

GLIC (***G****loeobacter* **l**igand-**g**ated **i**on **c**hannel) is a transmembrane protein of *Gloeobacter violaceus*, a unicellular cyanobacterium, and represents a prokaryotic member of the superfamily of pentameric ligand-gated ion channels (pLGICs) (1,2). Like other pLGICs, GLIC comprises of five protein subunits, and in its open conformation is arranged in a funnel-shaped structure (3,4). Each subunit contains a ligand-binding extracellular domain (ECD) and a pore-forming transmembrane domain (TMD). Most of the ECD encompasses a β-sandwich consisting of two antiparallel β-sheets. The TMD of a GLIC subunit contains four α-helices (M1, M2, M3, and M4). This overall fold is conserved across prokaryotic and eukaryotic pLGICs, such as serotonin type 3A (5-HT_3A_), nicotinic acetylcholine (nACh), and GABA_A_ receptors (Fig. 1). These receptors are implicated in diverse diseases and disorders of the human central nervous system, such as Alzheimer’s disease, schizophrenia, autism spectrum disorder, anxiety, attention-deficit/hyperactivity disorder, and premenstrual dysphoric disorder (PMDD) (5-11).

**FIGURE 1.**
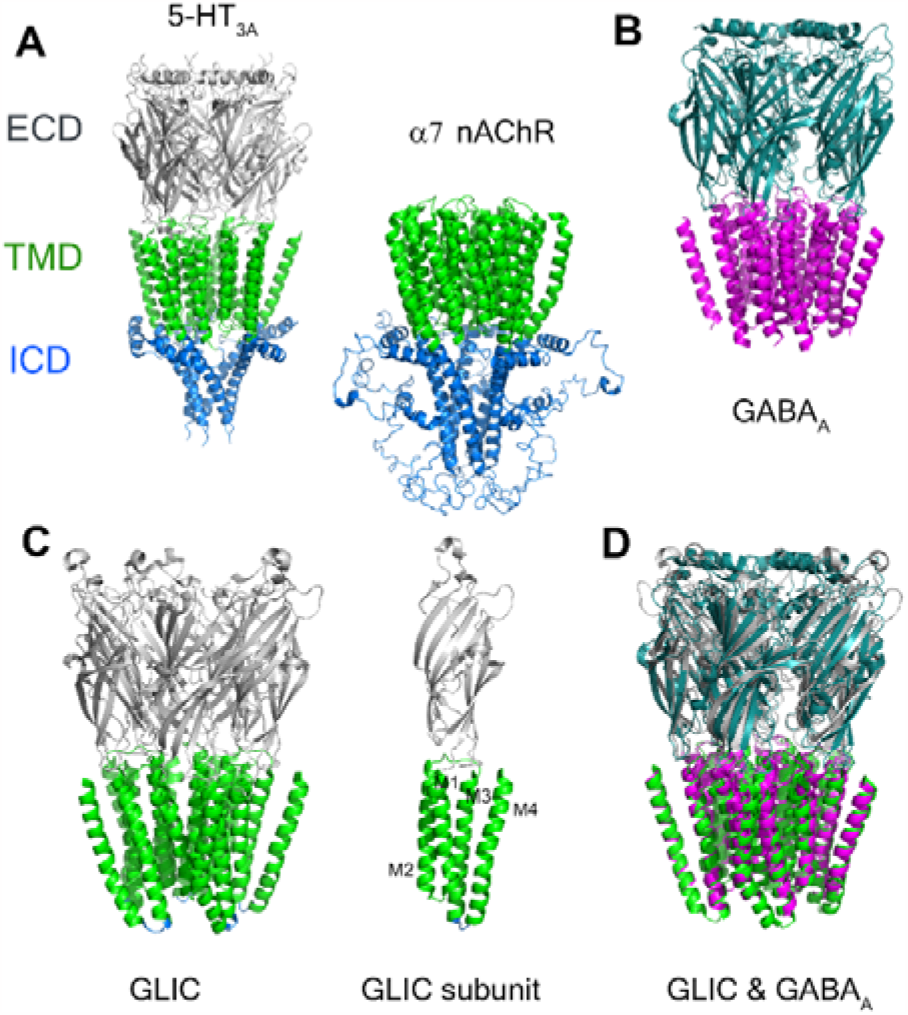
Ribbon representation of GLIC and other pentameric ligand-gated ion channels viewed parralel to the membrane plane. **A**, Crystal structure of serotonin type 3A (5-HT_3A_, PDB ID: 4PIR) and NMR structure of alpha 7 nicotinic acetylcholine receptor sub-type without ECD (α7 nAChR, PDB ID: 7RPM). ECD - extracellular domain. TMD - transmembrane domain. ICD-intracellular domain. **B**, Cryo-EM structure of GABA_A_ receptor (PDB ID: 7QN6). **C**, Crystal structure of *Gloeobacter violaceus* ligand-gated ion channel (GLIC, PDB ID: 6HZW); M1-M4 are transmembrane segments 1, 2, 3, and 4. **D**, Structural Alignment of GLIC (gray and green) and GABA_A_ receptor (marine and magenta).

In contrast to eukaryotic pLGICs that have an intracellular domain (ICD) of variable lengths between ca. 50 to 280 amino acids, the M3-M4 linker in GLIC is just four amino acids long (Fig. 1 A and C).

Because of this absence of an ICD and the remarkable structural homology of GLIC to other pLGICs (Fig. 1 D), GLIC has been used as a structural and functional surrogate model to understand the inhibition and binding mechanisms of antidepressants, anesthetics, cholesterol, and neurosteroids, the modulators of eukaryotic pLGICs such as 5-HT_3A_, nACh, and GABA_A_ receptors (12-17). Several binding pockets for anesthetics and neurosteroids, such as propofol, diazepam, and allopregnanolone in GABA_A_ receptors, are presented in Fig. 2 (16,18-23).

**FIGURE 2.**
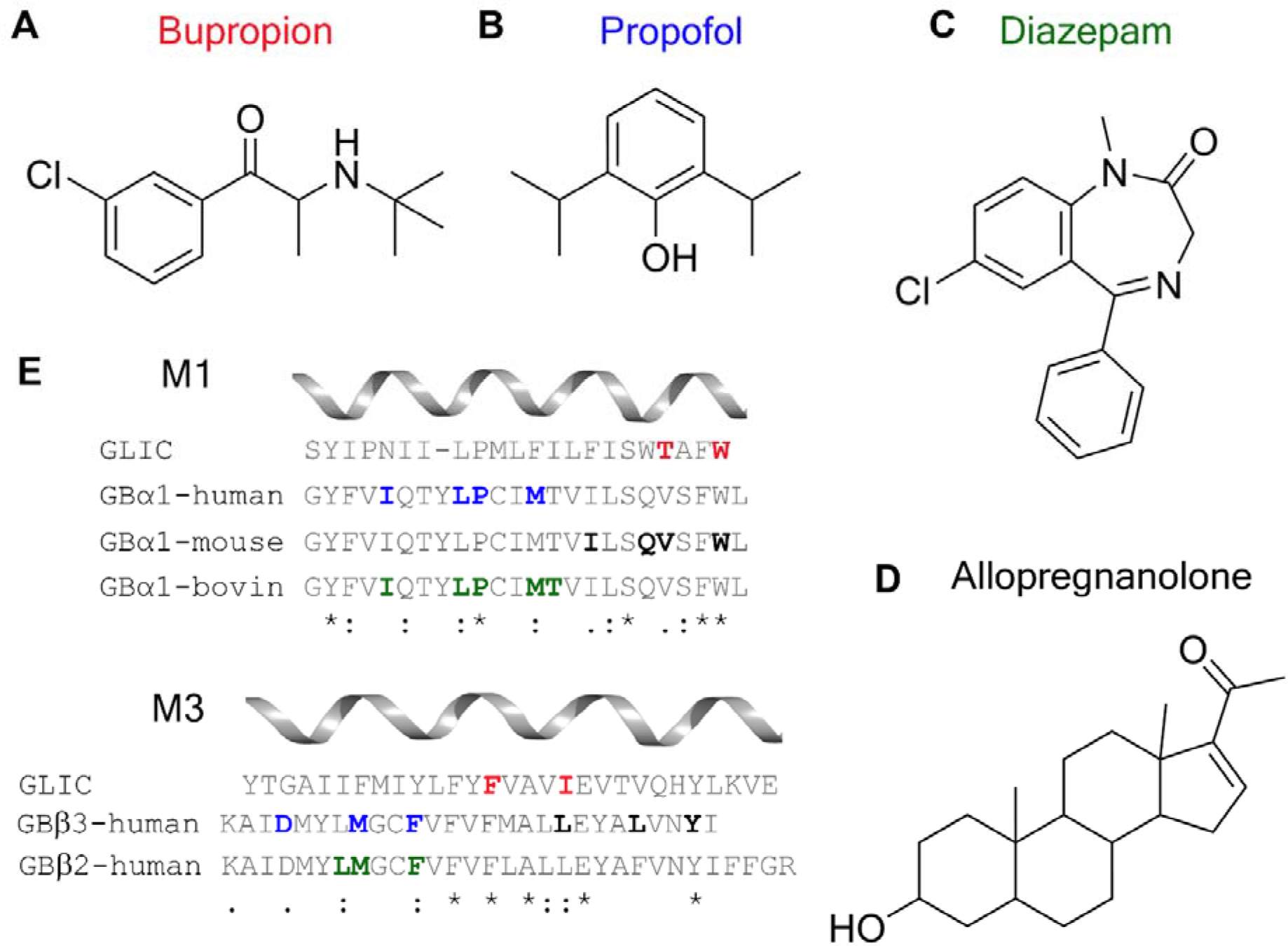
Binding site residues within the TMD for modulators of GABA_A_ receptors. **A-D**, Representatives of drug classes that modulate other pentameric ligand-gated ion channels and GABA_A_ receptors. **E**, Alignment of amino acid residues in M1 and M3 segments of TMD domains from GLIC and subunits of GABA_A_ receptors. Bupropion, an atypical antidepressant, binds to other pLGICs, such as serotonin type 3A receptors and nicotinic acetylcholine receptors; in this work, residues in red of M1 and M3 GLIC were mutated to examine the binding characteristics of bupropion in GLIC. Residues contributing to binding sites for propofol, an anesthetic, are in blue in M1 and M3 segments. The binding pocket for diazepam, a benzodiazepine, includes residues in green. Allopregnanolone, a neurosteroid, binds to residues in bold black.

As a proton-gated cation-selective channel, GLIC is activated by a pH decrease on the extracellular side (2). When activated at a low pH, a conformational change causes the channel to open, permitting positive charged ion influx through the central ion channel pore that is lined by the five M2 helices of the TMDs within the pentameric receptor.

Bupropion, Fig. 2 A, is a prescription medication used to treat depression and attention-deficit/hyperactivity disorder, to support smoking cessation, and recently to manage negative symptoms of schizophrenia (24-26). However, at high doses and in chronic treatments, bupropion and hydroxybupropion can cause many adverse effects, such as irritation, aggressive behaviour, paranoid delusion, and cardiogenic shock (27-29). When used at lower doses, other adverse effects of bupropion have also been reported, such as insomnia, anxiety, confusion, and tremors (30). Previous studies show that bupropion inhibits the uptake of dopamine and norepinephrine (31,32). This compound and its metabolites are also non-competitive inhibitors of many pLGICs, such as nicotine acetylcholine receptor subtypes α4β2, α3β4, and α7, and serotonin type 3 receptors (33-37). However, the bupropion binding sites on these channels are still elusive.

To further our knowledge of the mechanism of bupropion action on pLGICs, we examined the impact of single amino acid substitutions in GLIC on the bupropion inhibitory potency. We introduced single amino acid substitutions at residues T214 and W217 in the M1 transmembrane segment and residues F267 and I271 in the M3 transmembrane segment (Fig. 2 E). GLIC constructs containing individual substitutions, one at a time, were then expressed in *Xenopus* oocytes. The proton-induced dose-response relationship and the inhibitory effect of bupropion in these GLIC constructs were measured using two-electrode voltage-clamp recordings and analyzed using one-way ANOVA with Tukey’s post-hoc test. We determine that substitutions W217C and F267W shift the pH_50_ of GLIC 0.9- and 1.3-fold, respectively. While the inhibitory potency of bupropion for constructs T214C, W217C, and I271W was comparable with wildtype, substitution T214F induced a 13-fold shift in IC_50_. We thus infer that T214 is involved in shaping the binding site for bupropion in GLIC.

## MATERIALS AND METHODS

### Construct Design

The GLIC coding region from *Gloeobacter violaceus* (ATCC 29082) was cloned into the pXOON vector as described previously (38). Single amino acid substitutions of GLIC were engineered using the QuikChange II Site-Directed Mutagenesis kit (Agilent Technologies) and oligonucleotide primers (Sigma). The DNA sequence of the entire coding region was then verified by Genewiz/Azenta to obtain point mutations T214F, W217C, F267W, T214C, I271W, or T214A. Amino acid positions are numbers as in crystal structure with PDB ID of 6HZW(39).

### Expression of GLIC channels in *Xenopus laevis* oocytes

Plasmid constructs were linearized using *Xba*I restriction enzyme (New England Biolabs) to create DNA templates for complementary RNA (cRNA) production. cRNAs were produced using the mMESSAGE mMACHINE T7 kit (Applied Biosystems/Ambion). The capped mRNAs were then purified using the MEGAclear kit (Applied Biosystem/Ambion), dissolved in nuclease-free water, and stored at -80°C until use. Defolliculated *Xenopus* oocytes were purchased from Ecocyte Bioscience US LLC. Oocytes were injected in OR2 buffer (82.5 mM NaCl, 2 mM KCl, 1 MgCl2 mM, 5 HEPES, pH 7.5) and then kept in OR2 buffer containing 5% horse serum (Sigma-Aldrich, Cat. # 26050088) and 1x antibiotic-antimycotic (ThermoFisher Scientific/Gibco Cat. # 15240-062) at 15°C. Oocytes were injected with 50.6 nl of 200 ng/μl cRNA (10 ng) for expression of the individual wild type and engineered GLIC channels. After the injection, the oocytes were incubated at 15°C for 2-4 days before being used for two-electrode voltage-clamp recordings.

### Electrophysiology

Electrophysiological recordings of GLIC ion channels were obtained via the two-electrode voltage clamp (TEVC) method. Each oocyte was placed in a 200 μl perfusion chamber containing the GLIC oocyte recording buffer, GORB, pH 7.5 (100 mM NaCl, 20 mM NaOH, 2.5 mM KCl, 2 mM CaCl_2_, 1 mM MgCl_2_, 5 mM citric acid, 5 mM HEPES), which was perfused into the chamber at a rate of 5-6 ml/min. The ground electrode containing 1% agarose in 3 M KCl was connected to a bath shaping an extension of the perfusion chamber. The voltage and current electrodes were filled with 3M KCl solution with resistances in the range of 0.5 to 2 MΩ. Each oocyte was voltage-clamped at a holding potential of -60 mV in a setup with a TEV-200A Voltage Clamp amplifier (Dagan Corporation), and data were acquired using a Digidata 1440A digitizer (Molecular Devices) and Clampex 10.7 software with gap-free acquisition mode and 200 Hz sampling rate. Different proton concentrations ranging from pH 4.0 to pH 7.5 were used to elicit currents of GLIC to determine the pH at which 50% of the maximal response or maximal current amplitude was recorded, pH_50_.

### Determining pH_50_ for GLIC

Currents were recorded for individual oocytes with increasing proton concentrations or decreased pH values. The GLIC oocyte recording buffer (GORB) was varied in pH starting at the pH of 7.5 and decreased down to a pH of 4 or 3.7 at a step interval of about 0.5 pH units. Perfusion with GORB of pH 7.5 provides a resting current for comparison, which equals zero, where the channels are closed. After recording GLIC currents with GORB of the lowest pH (pH 4 or 3.7), the oocyte was superfused with GORB, pH 7.5, to determine the complete reversibility of low pH induced currents. Currents recorded from unstable oocytes that did not return to the resting current at pH 7.5 were not included in the data set for analysis.

The current (*I*) elicited at a given pH obtained by subtracting the peak current from the resting current was normalized to the maximum current amplitude (*I*_*max*_) obtained from the same oocyte in the same set of tested pHs. Data points obtained from individual oocytes were then fitted using nonlinear regression in GraphPad Prism 9 (log(agonist) vs. normalized response – variable slope). A nonlinear proton concentration-current relationship was built, and LogEC_50_ and Hill coefficient (n_H_) were determined using the below Hill equation.

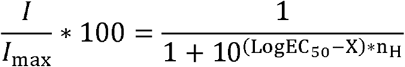

*I*_*max*_ is the current activated at the highest proton concentration used in a set of tested pHs; EC_50_ is the proton concentration at which 50% of *I*_*max*_ is achieved; Negative LogEC_50_ (-LogEC_50_) is the pH_50_; X is the logarithm of a proton concentration used in the tested pHs; and n_H_ is Hill coefficient that describes the steepness of the dose-response curve. When n_H_ is greater than -1, the dose-response curve is shallower than the standard curve. When n_H_ is smaller than -1, the dose-response curve is steeper than the standard curve.

The averages or means of pH_50_ and n_H_ values were obtained from at least 3 sets of tested pHs or n ≥ 3 oocytes. The statistical significance was determined using one-way ANOVA with Tukey’s post-hoc test. Data are represented as the mean ± SEM.

### Determining bupropion IC_50_ for GLIC

TEVC recordings were conducted as described above, where bupropion was dissolved in the buffer of the pH corresponding to the respective pH_50_. Bupropion was titrated with increasing concentrations from 0 μM to 1000 μM to determine a bupropion concentration producing 50% inhibition of *I*_max_ (IC_50_). *I*_max_ is the current produced at the pH_50_ without bupropion. To fit an inhibition-response curve from a set of bupropion concentrations, the nonlinear regression model in GraphPad Prism 9 (inhibitor vs. normalized response – variable slope) was used. The IC_50_ and Hill coefficient values were determined using the below Hill equation, where *I* is an inhibited current produced at a bupropion concentration. X is the logarithm of a bupropion concentration used in the experiment, and n_H_ is the Hill coefficient describing the dose-response curves’ steepness.

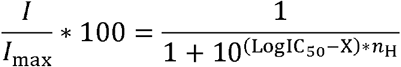

The averages or means of IC_50_ and n_H_ were obtained from 3 sets of bupropion concentrations collected from n = 3 oocytes. The statistical significance was determined using one-way ANOVA with Tukey’s post-hoc test. Data are represented as the mean ± SEM.

All figures and graphs were made using GraphPad Prism 9, Adobe Illustrator CC 2023, and Adobe Photoshop 2023.

## RESULTS

Agonist binding sites in pLGICs have been identified at intersubunit interfaces of the ECD many years ago (40-46). Additionally, an abundance of allosteric modulator sites has been described at intersubunit interfaces in both the ECD and TMD. Small ligands binding at these sites that bring adjacent subunits into closer apposition promote concerted motions and act as agonists, whereas compounds that destabilize intersubunit interactions act as antagonists (47). In GABA_A_ receptors, the interfaces between M1 and M3 of adjacent subunits house binding sites for many clinically used drugs such as propofol, diazepam, and allopregnanolone (Fig. 2) (18,19,48). Based on the lipophilicity of bupropion and preliminary molecular docking screening for GLIC and antidepressants, residues T214 and W217 in M1 and residues F267 and I271 in M3 of the GLIC transmembrane domain were suggested to participate in bupropion modulation of GLIC (data not shown). We generated eight GLIC constructs, each containing a single amino acid substitution at either of these positions T214, W217, F267, or I271 (Fig. 2 E and Fig. 3 A). We then expressed these constructs and wild-type GLIC in *Xenopus laevis* oocytes. After 3-5 days of protein expression, two-electrode voltage-clamp recordings were used to measure proton-induced currents at different proton concentrations or pH values, and in addition bupropion-inhibited responses at the respective pH_50_ of individual GLIC constructs. Effects of single amino acid substitutions on these responses and potential binding sites for bupropion in GLIC were then determined, as described below.

**FIGURE 3.**
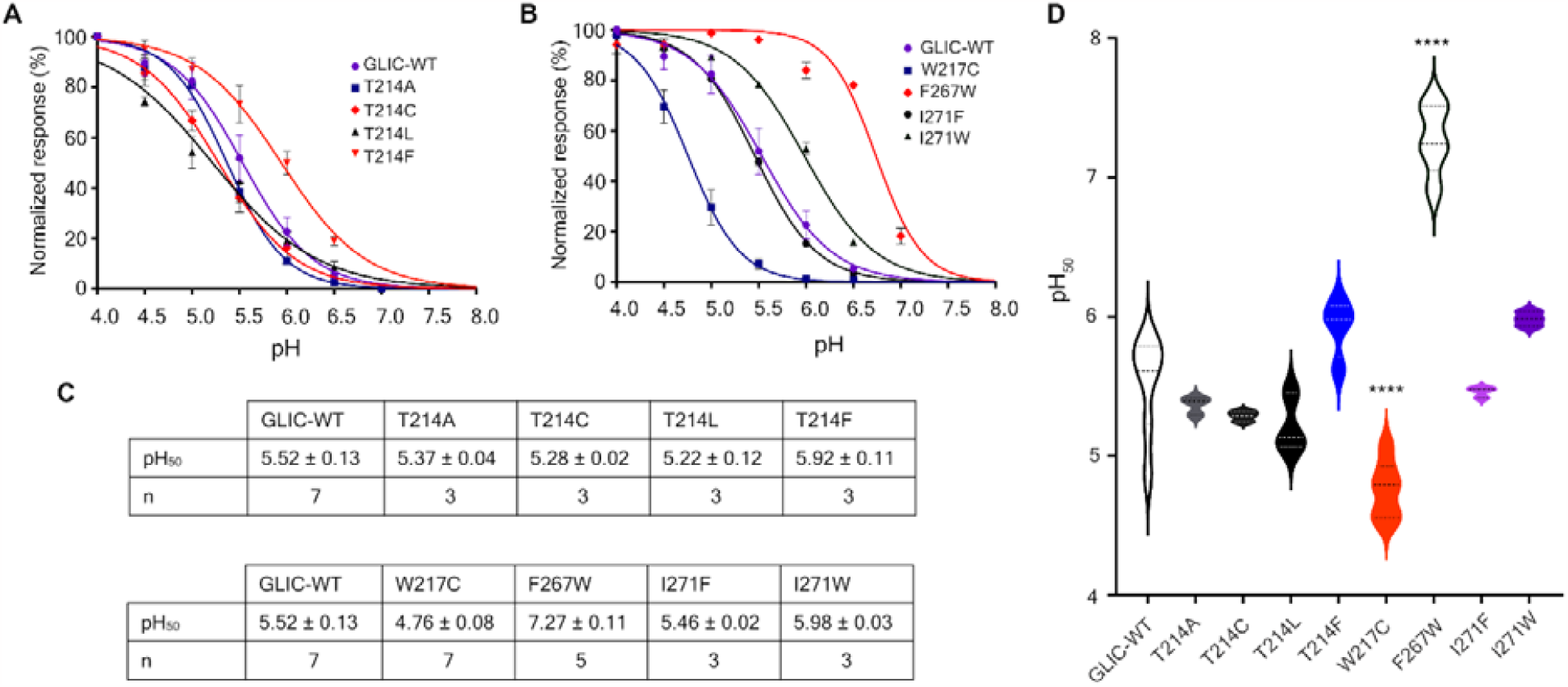
Responses of GLIC constructs at different pH values. **A**, pH-response curves of currents elicited from wild-type GLIC (WT) and substitution constructs. Single amino acid substitutions were made at residue T214. **B**, pH-response curves of currents elicited from wild-type GLIC (WT) and GLIC constructs. The single substitutions were made at residues W217, F267, and I271. Mutation W217C shifts the pH50 to more acidic pH, while mutation F267W shifts the pH50 to more basic pH. **C**, Negative LogEC_50_ or pH_50_ and the number of biological replicates (n) for different GLIC constructs. pH_50_ is the pH value at which the current amplitude is halfmaximal. **C**, Significant differences in pH_50_ were determined using one-way ANOVA with Tukey’s post-hoc test. Significance is indicated as **** in W217C with p= 0.00002 and **** in F267W with p = 0.0000000000031. Data are shown pH_50_ ± SEM and Hill coefficient ± SEM from n ≥ 3 oocytes.

### Effects of mutations in M1 and M3 segments of GLIC transmembrane domain on proton-activated currents

To examine the effect of mutations on the pH-gated property of GLIC, we measured proton-induced responses of the GLIC constructs by applying increasing proton concentrations as described in the methods section. The current amplitudes increased from higher to lower pH values, as seen in the proton-response curves and the current traces (Fig. 3 A and B, Fig. 4). We determined the proton concentration at which the response was half-maximal, pH_50_, for each construct.

**FIGURE 4.**
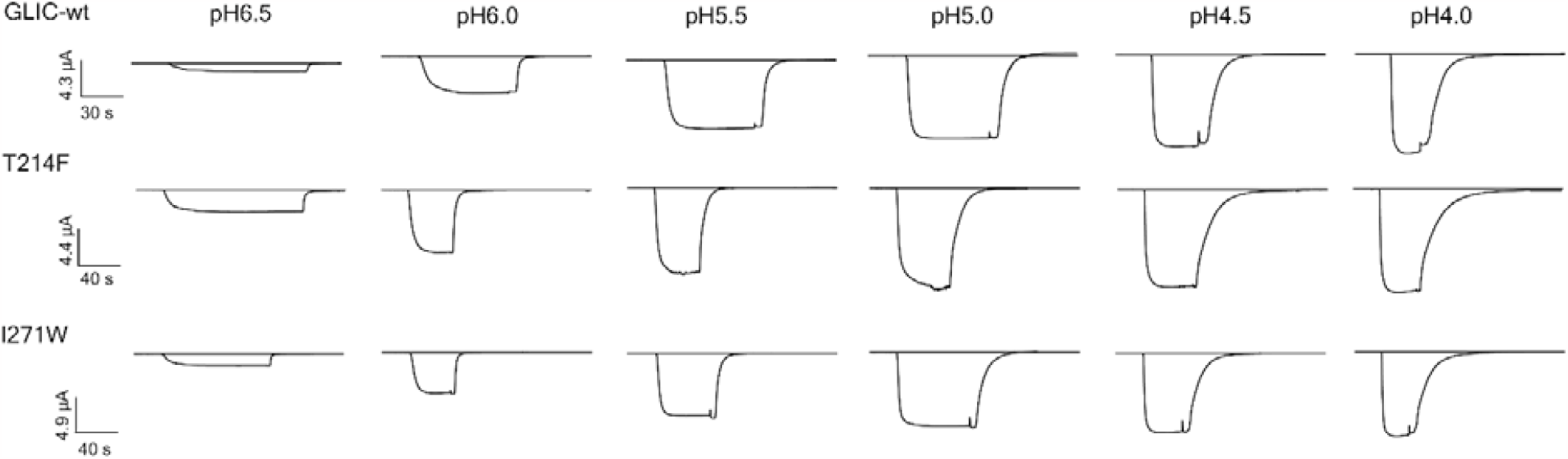
Representative current traces of the wild-type GLIC (GLIC-wt), T213F GLIC, and I270W GLIC evoked at different pHs. Current amplitudes are smaller at higher pHs.

The wild-type GLIC gave rise to a pH_50_ of 5.52 ± 0.13 with a Hill coefficient n_H_ of -1.41 ± 0.08, Fig. 3 C, consistent with the typical pH-gated features of this prokaryotic channel described previously (2,38,49).

For single amino acid substitutions at residue T214, such as T214A, T214C, T214L, and T214F, the pH_50_ values of 5.37 ± 0.04, 5.28 ± 0.02, 5.22 ± 0.12, and 5.92 ± 0.11, respectively, were determined (Fig. 3 D, top panel). No significant difference between pH_50_ values of either of these constructs vs. wild-type GLIC was observed (Fig. 3 E). We also observed no significant differences in Hill coefficients for these constructs. The amino acid substitutions introduced at residue T214 in M1 of the GLIC transmembrane domain, thus, do not alter proton-induced current responses of GLIC. We also observed no significant difference in pH_50_ values or Hill slopes for I271F and I271W GLIC constructs, with pH_50_ values of 5.46 ± 0.02 and 5.98 ± 0.03 (Fig. 3 C, lower panel, Fig. 3 D).

However, a statistically significant difference in pH_50_ values was seen for wild-type GLIC vs. either W217C GLIC or F267W GLIC construct, with a p = 0.00002, or p = 0.0000000000031, respectively, using one-way ANOVA with Tukey’s post-hoc test (Fig. 3 C, lower panel and Fig. 3 D). Substitution W217C shifted the pH_50_ of to a more acidic pH compared to wild type, from 5.52 ± 0.13 down to 4.76 ± 0.08. In contrast, mutation F257W shifted the pH_50_ to a more basic value, from 5.52 ± 0.13 up to 7.27 ± 0.11. Our data here suggest that GLIC with a Cys residue at position 217 in M1 requires a higher proton concentration for opening, while GLIC with a Trp residue at position 257 in M3 requires less protons for opening. No significant difference was observed in Hill coefficients of either the W217C GLIC or F267W GLIC construct vs. the wild-type GLIC. These changes in sidechains at residue W217 in M1 and residue F267 in M3 of the GLIC transmembrane domain, thus, allosterically modulate channel opening.

### Bupropion modulation and binding sites in GLIC

To identify potential bupropion binding sites in the M1 and M3 segments of the GLIC TMD, we next measured bupropion inhibition response curves. We then calculated the IC_50_ and Hill coefficient values for wild-type, T214C, T214F, W217C, and I271W GLIC constructs as described in the methods section. Residue T214, W217, and I271 reside at the interface of two adjacent subunits of GLIC as illustrated in Fig. 5 A.

**FIGURE 5.**
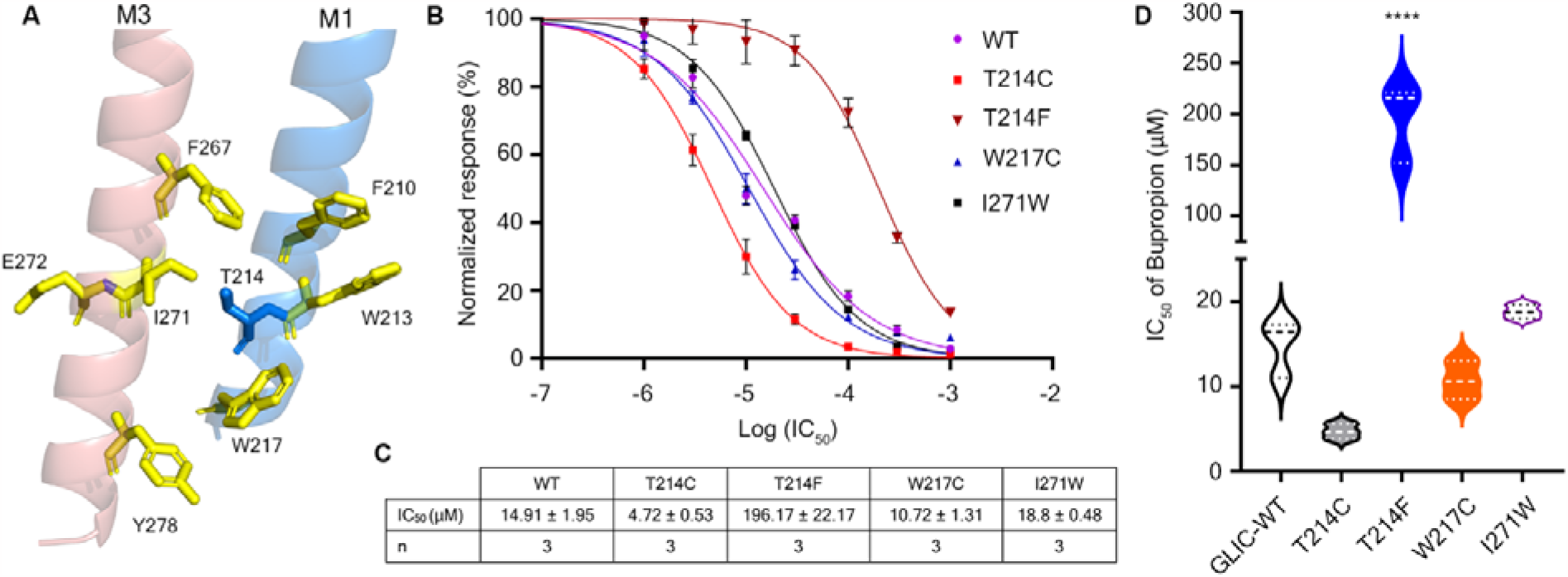
Inhibition of proton-inducted currents with bupropion. **A**, Location of residues T214, W217, and I271 at the intersubunit binding pocket for neurosteroid, allopregnanolone, in GLIC – residues contributes to this pocket are highlighted in yellow stick (16). **B**, Dose response curves for proton-gated GLIC currents with increasing concentrations of bupropion. **D-C**, IC_50_ values of bupropion for different GLIC constructs. Data are shown IC_50_ ± SEM from n = 3 oocytes. Significant differences in responding to bupropion concentration were observed for the wild-type GLIC vs. the T214F GLIC (one-way ANOVA with Tukey’s post-hoc test, **** with p = 0.000001).

We observed that bupropion inhibited the proton-induced currents of wild-type GLIC with an IC_50_ of 14.91 ± 1.95 μM (Fig 5 B and C). Mutations T214C and W217C produced smaller IC_50_ values, 4.72 ± 0.53 μM, and 10.72 ± 1.31 μM, respectively, while mutation I271W produced a larger IC_50_ of 18.8 ± 0.48 μM (Fig 5 A and B); however, these differences in the IC_50_ values are not statistically significant compared to the wild-type in our study. Previous work by Cheng et al. on mutation I271W indicates this mutation does not affect the inhibition of allopregnanolone; however, I271W significantly reduces the inhibition of KK200, a neurosteroid analogue (16). The counterpart of this Ile residue in the M3 segment of GABA β2 TMD, residue L297, lines the binding pocket of allopregnanolone as recently reported by Gouaux and co-workers(19) (Fig. 2 E).

While replacing an ethyl hydroxyl side-chain with a methyl thiol side-chain at position 214 in M1 (T214C) did not affect the bupropion inhibitory effect in GLIC, adding a methylphenyl side-chain at this position (T214F) dramatically decreased the potency of bupropion toward GLIC. The IC_50_ of T214F, 196.17 ± 22.17 μM, was reduced 13-fold compared to wild-type, IC_50_ of 14.91 ± 1.95 μM (Fig. 5 C and D). We infer that the substitution T214F impairs or interferes with the binding and consequently the inhibitory effect of bupropion to GLIC. Our data, thus, indicates that residue T214 lines the binding pocket for bupropion in GLIC.

## DISCUSSION

In this work, we examined the effects of single amino acid substitutions in the lower third of transmembrane segments M1 and M3 in GLIC on pH-gating and the bupropion potency. We find that mutation W217C shifts the pH_50_ of GLIC to a more acidic pH, and mutation F267W shifts the pH_50_ to a more basic pH, respectively. We also find that bupropion inhibits GLIC, and that substitution of residue T214 in the M1 segment with a Phe residue reduces bupropion inhibitory activity and therefore T214 likely lines the binding pocket for bupropion in GLIC.

By shifting the response of GLIC toward a higher proton concentration compared with the wild type (Fig. 3), mutation W217C might help to promote an open state. In contrast, mutation F267W might help to stabilize the closed state of GLIC when shifting the response of GLIC toward a lower proton concentration. In other words, residues W217 and F267 play critical roles in regulating GLIC conformational changes upon the interaction of the channel with its agonist, protons. Both tested residues Phe and Trp are conserved in TMD M1 and M3 of GLIC and GABA_A_ receptors but not in other eukaryotic subunits of pLGICs. Residue W217 contributes to the intersubunit binding pocket of allopregnanolone (16). And when Cheng and co-workers substituted Trp with an Ala residue at this position, W217A GLIC construct does not produce functional receptors. The W217 counterpart residue in GABA_A_ receptors is W245 in M1 of the α1 subunit, and this residue W245 lines the binding sites of the neurosteroids (Fig. 2 E) (19,21,23). This conserved Trp residue in M1 thus contributes to the allosteric modulation either directly via its lipophilic side chain as observed by the change in pH_50_ and/or via binding of lipophilic allosteric modulators like neurosteroids.

Bupropion is an atypical antidepressant and smoking cessation drug, which causes many adverse effects, and the most common ones are insomnia, irritability, and anxiety (30). Whether or not these adverse effects result from the interactions of bupropion with neurotransmitter-gated ion channels or pLGICs such as GABA_A_, serotonin type 3A, or neuronal nACh receptors is still under investigation. Bupropion inhibits nicotinic acetylcholine receptors in a non-competitive manner (33,34,50). Our previous work also determined that bupropion and its metabolite, hydroxybupropion, block the function of serotonin type 3A receptors within its therapeutically-relevant concentrations (36,37).

In this work, we report the inhibitory potency of bupropion for a prokaryotic member of the pentameric ligand-gated ion channels, GLIC, with an IC_50_ of 14.91 ± 1.95 μM (n=3, Fig 5 B and C), which is similar to previously observed for bupropion potency in eukaryotic pLGICs (34,36,37,50). Additionally, we show that the bupropion potency is dramatically reduced by the T214F substitution but not by T214C (Fig. 5 B-D). We infer that the longer side chain of the Phe residue leads to a direct steric clash with bupropion, leading to a higher concentration of bupropion being required to inhibit the channel. The corresponding residues to GLIC T214 in other subunits of pLGICs, which are either Leu or Val residues, possess longer side chains as compared to the Thr residue. Perhaps so, the IC_50_ of bupropion is also comparatively higher in serotonin type 3A receptors (87 μM), serotonin type 3AB receptors (840 μM), α7 nicotinic acetylcholine receptors (54 μM), and α4β2 nicotinic acetylcholine receptors (17.8 μM) (34,36,37,50). We infer that T214 directly lines the bupropion binding site. Residue T214 is located within the intersubunit binding pocket of neurosteroids in GLIC, Fig 5 A (16). The corresponding residue to T214, residue V242 of the α1 subunit of murine GABA_A_ receptors, has also been observed to line the binding pocket for neurosteroids (Fig. 2 E) (19). A nearby residue, W217 in GLIC, and its counterpart, residue W245 of the α1 subunit of GABA_A_ receptors, also contributes to the neurosteroid binding pocket (16,19,21,23). Taken together, bupropion when bound to this site in GLIC would extend from T214 to W217, yielding an overall comparable binding pocket to neurosteroids in pentameric ligand gated ion channels.

## AUTHOR CONTRIBUTIONS

EP, AP, and ZRG recorded the data. EP, AP, ZRG, GF, and HQD analyzed the data. HQD and MJ wrote the manuscript with the other authors contributing draft sections. All authors contributed to the final version of the manuscript.

## ACKNOWLEDGMENT

We thank Dubem Onyejegbu, Jessica Shepherd, and Emma Brackett for support with some of the data collected. Research reported in this publication was supported by the National Institute of Neurological Disorders and Stroke of the National Institutes of Health under award number R01NS077114 (to M.J.).

